# Signatures of negative frequency dependent selection in colonisation factors and the evolution of a multi-drug resistant lineage of *Escherichia coli*

**DOI:** 10.1101/400374

**Authors:** Alan McNally, Teemu Kallonen, Christopher Connor, Khalil Abudahab, David M. Aanensen, Carolyne Horner, Sharon J. Peacock, Julian Parkhill, Nicholas J. Croucher, Jukka Corander

**Author notes:** Equal contributions (Alan McNally, Teemu Kallonen, Christopher Connor). Equal contributions (Nicholas J. Croucher, Jukka Corander).

## Abstract

*Escherichia coli* is a major cause of bloodstream and urinary tract infections globally. The wide dissemination of multi-drug resistant (MDR) strains of extra-intestinal pathogenic *E. coli* (ExPEC) poses a rapidly increasing public health burden due to narrowed treatment options and increased risk of failure to clear an infection. Here, we present a detailed population genomic analysis of the ExPEC ST131 clone, in which we seek explanations for its success as an emerging pathogenic strain beyond the acquisition of antimicrobial resistance (AMR) genes. We show evidence for evolution towards separate ecological niches for the main clades of ST131 and differential evolution of anaerobic metabolism, key colonisation and virulence factors. We further demonstrate that negative frequency-dependent selection acting across accessory loci is a major mechanism that has shaped the population evolution of this pathogen.

## Introduction

*Escherichia coli* is now the most common cause of blood stream infections in the developed world, outnumbering cases of *Staphylococcus aureus* bacteraemia by 2:1 ^1^. *E. coli* is also the most common cause of urinary tract infections (UTI), which in turn are among the most common bacterial infections in the world ^2^. Bacteraemia and UTI are caused by a subset of *E. coli* termed extra-intestinal pathogenic *E. coli* (ExPEC). ExPEC are not a phylogenetically distinct group of *E. coli* but rather represent strains which have acquired virulence-associated genes that confer the ability to invade and cause disease in extra-intestinal sites ^3^. Genes associated with virulence that confer the ability to adhere to extra-intestinal tissues, to sequester extracellular iron, to evade the non-specific immune response, and toxins resulting in localised tissue destruction have all been described as essential in the process of ExPEC pathogenesis ^4^.

The problem presented by the scale of ExPEC infections is exacerbated by the number of cases involving multi-drug resistant (MDR) strains ^1,5,6^. Epidemiological surveys report as many as 60% of UTI ExPEC isolates as being resistant to three or more classes of antibiotics, and as many as 50% of bacteraemia isolates ^5,6^. The increase in MDR ExPEC prevalence has been rapid and primarily attributable to a small number of ExPEC lineages ^5^. The most common of these is the *E. coli* ST131 lineage, which has rapidly become a dominant cause of ExPEC UTI and bacteraemia globally ^5–7^. *E. coli* ST131 is particularly associated with carriage of the CTX-M class of extended-spectrum β-lactamase (ESBL) which confers resistance to 3^rd^-generation cephalosporins ^7^, and there have been a small number of reports of *E. coli* ST131 isolates carrying metallo-β-lactamases conferring resistance to carbapenems ^8^. The carriage of these resistance genes is driven by acquisition and stable maintenance of large MDR plasmids ^9^.

The phylogenetic structure of *E. coli* ST131 is well characterised ^10–14^ and shows the emergence of a globally disseminated, MDR-associated clade C from primarily drug susceptible clades A and B. The lack of phylogeographic signal and phylogenetic structure based on host source suggests rapid global dispersal and frequent host transitions within clade C ^14^. Research has suggested that the acquisition of fluoroquinolone resistance via point mutations in DNA gyrase and DNA topoisomerase genes was the primary driver in the rapid emergence of clade C, alongside the predated acquisition of well-defined ExPEC virulence factors ^11,12^. Later work also suggested that clade C *E. coli* ST131 may dominate as a successful MDR clade due to the ability to offset the fitness cost of MDR plasmid acquisition and maintenance via compensatory mutations in gene regulatory regions ^14^. Genome-wide association studies (GWAS) have been used to identify loci and lineage specific alleles significantly associated with clade C *E.coli* ST131, which suggested a secondary flagella locus encoding lateral flagella (Flag-2^15^), and a number of hypothetical proteins and promoter regions as being clade C *E. coli* ST131 associated loci ^14^.

Recent work on *E. coli* causing bacteraemia provided compelling evidence that resistance to antimicrobials has not been the major driver of the success of ST131 ^16^. Analysis of a large 11-year population survey across the UK showed that ST131 rapidly stabilised at a level of approximately 20% after its emergence around 2002 in the UK. This was far in excess of already-resident MDR clones, such as ST88 or ST405. Nevertheless, the overall prevalence of resistance phenotypes remained approximately constant in the population. Furthermore, most currently known major ExPEC clones (primarily ST12, ST73, ST95, and ST69, the last of which also rapidly emerged in 2002) show a similar stable population frequency across the 10 years following the introduction of ST131, despite exhibiting far less extensive resistance profiles. These observations suggested the distribution of ExPEC strains was shaped by negative frequency-dependent selection (NFDS) ^16^. NFDS describes the situation in which a given phenotype is most beneficial to a population when it is rare. This is because as the phenotype becomes common it becomes costly, because of pressures such as host response to the population.

Recently a multilocus NFDS model of post-vaccination *Streptococcus pneumoniae* population dynamics has been described ^17^. Frequencies of accessory genes were found to be highly conserved across multiple populations on different continents, despite these populations themselves being composed of different strains, as defined by core genome sequences. Detailed modelling and functional analysis indicated changes in strain prevalence could be explained by NFDS driving accessory loci towards equilibrium frequencies, through mechanisms involving interactions with other bacteria, hosts, or mobile elements ^17^. The level of the selective force was estimated to be similar across the populations and manifested itself in the maintenance of stable population frequencies of accessory loci, despite a substantial perturbation of the population by the introduction of the pneumococcal vaccine ^17^.

Here, we present the analysis of 862 genomes collated from previous large scale *E. coli* ST131 phylogenomic studies ^11–14,16,18^ and newly sequenced isolates from the BSAC bacteraemia resistance project from the UK and Ireland. This allowed us to perform sufficiently powered population genetic analyses and identify the key steps in the evolution from the largely drug susceptible clades A and B to the globally dominant MDR clade C. Pan-genome analyses identified the formation of clade C was underpinned by an accumulation of allelic diversity, particularly enriched for genes involved in anaerobic metabolism and other loci important for colonisation of the human host by ExPEC. Our data suggest the evolution of the MDR phenotype is part of a wider, ongoing adaptation towards prolonged human colonisation that is currently accompanied by a radiation through diversification of metabolic and antigenic loci.

## Methods

### Genome data

We analysed a collection of 862 *E. coli* ST131 genomes (Table S1), of which 684 were previously sequenced as part of phylogenomic investigations of the ST131 lineage ^10,11,13,14,16,19^. We added 184 previously unpublished ST131 isolates from the British society for antimicrobial chemotherapy (BSAC) bacteraemia resistance surveillance project which were selected from *E. coli* in the BSAC resistance surveillance bacteraemia collection from the UK and Ireland between 2001–2011. Together this collection represents bacteria isolated from invasive disease (blood stream infections), human asymptomatic carriage and disease resulting from intestinal carriage (UTI), and bacteria isolated from a range of veterinary livestock, pets, wild birds, and the wider environment, minimising population or sampling bias to as large an extent as possible.

In an attempt to avoid any issues arising from different assembly or annotation metrics employed in the previous projects, we downloaded only raw sequence data in fastq format using the previously published accession data. We then performed de novo assembly on all the genomes using Velvet ^20^ and annotation using Prokka ^21^ as previously described ^16^. A pan-genome of the entire data set was constructed using Roary with 95% identity cut-off ^22^. A concatenated core CDS alignment was made from the Roary output and a maximum likelihood phylogenetic tree was constructed from the alignment using RaxML version 8.2.8 ^23^ and the GTR model with Gamma rate heterogeneity.

For comparative lineage analysis we utilised the 264 ST73 genomes, and162 ST95 genomes that were sequenced and fully characterised as part of the UK BSAC genome study ^16^.

### Accessory genome analysis

The pan-genome matrix from Roary was utilised to investigate the presence of clade specific loci. The PANINI tool was used with the default setting to visualize the accessory gene sharing patterns in the population https://microreact.org/project/BJKoeBt2b ^24^. PANINI has been demonstrated to provide efficient complementary visual means to phylogenetic trees to accurately extract both distinct lineages present in a population-wide genomic dataset, and to highlight clusters within lineages, that are explained by rapidly occurring, homoplasic alterations, such as phage infection. Roary was run on the entire data set using the default 95% sequence identity threshold to cluster genes, allowing us to separate genes based on allelic as well functional differences. Based on a frequency distribution histogram (Figure S1), we assigned a locus as being clade specific if it occurred at a frequency > 95% in one clade and at < 5% in the other two clades. Loci identified as clade specific were functionally annotated by performing a tBlastn analysis of the nucleotide sequence of the loci against the NCBI non-redundant database.

### Functional categorisation of pangenomes

To assess the functional composition of the accessory pangenome we assigned Gene Ontology (GO) terms to gene sequences from the pangenome. Briefly, representative sequences from the pan genome of ST131 were mapped to orthologous groups in the bactNOG database using the eggNOG emapper utility ^25^ Mapping was performed using the diamond search algorithm. Output from eggNOG was filtered to remove Orthologous Groups with no GO terms, a score was assigned to each Orthologous Group based on gene mapping frequency.

### Comparisons of lineage and clade specific loci

In order to compare lineage pan-genomes whilst accounting for differences in the number of genomes a sampling approach was utilised. Specifically, a subset with size equal either to the number of ST73 or ST95 genomes was selected at random from the ST131 Clade C. The functional enrichment of genes in the subset was quantified and statistically compared to the ST73 or ST95 pangenome using a Chi Squared test. This process was repeated 100 times to produce 100 p-values, from which the median p-value was calculated. Utilising the same subsampling approach, the pangenome composition of Clade C ST131 genomes was compared to both the Clade A and Clade B pangenomes.

Chi squared statistical tests were performed to assess the significance of the observed differences in functional enrichment. Briefly, with each iteration of the sampling procedure a Chi squared test was performed using the functional proportion of the subsampled pangenome as the observed values and the proportions for ST73 or ST95 as the expected value. This generated 100 p-values from which one can use the average, maximum, or median to assess significance of the observed differences. In addition, proportional Z statistic tests were also performed to assess the significance of the observed difference. The measurements from the 100 replicates of the subsampling procedure were used to generate an average for the proportions as well as to estimate the variance. The tests were conducted using the proportional measurements from ST73 and ST95 as the ‘true’ means and quantifying how distinct the ST131 subsamples were from these reference values.

The sequences of 64 anaerobic metabolic genes in which allelic diversity was observed were extracted from individual genomes. The nucleotide sequences were then clustered at 80% identity and 80% length using CD-HIT which was run using the accurate flag and ‘word size’ of 5 ^26^. An additional CD-HIT script was used to extract gene sequences for clusters with more than 3 genes, the minimum required by MEGA-CC for analysis. The sequences were then aligned using Muscle with default settings ^27^. Resulting alignment files were analysed in MEGA-CC to produce measurements of Tajima’s D ^28^.

### ST131 clade specific SNPs

To visualise the ST131 clades A, B, C, C1 and C2 within the ML tree and the PANINI clustering we identified clade specific SNPs (Table S1) as previously described ^16^.

### NFDS modelling

NFDS modelling used genomic data from the previous publication analysing the population dynamic of blood stream infection *E. coli* isolates in the UK ^16^. Isolates were assigned to genotypes based on a hierBAPS analysis of the core genome ^29^. The previously-defined sequence types were used to divide any diverse clusters to similar levels of resolution. Therefore the clusters used corresponded to the largest hierBAPS cluster that corresponded to a clonal complex, if links were constructed between single- and double-locus variants; if neither condition could be satisfied, the third level of clustering was used. This identified 62 sequence clusters across the population. The sets of orthologous sequences were those defined by a previous Roary analysis ^16^ those present at between 5% and 95% frequency in the first sample, from 2001, were modelled as evolving under NFDS, and tending towards an equilibrium frequency, *e*_*l*_, corresponding to that in the 2001 sample.

Seven resistance phenotypes, present within this frequency range in 2001, were also modelled as evolving under NFDS: amoxicillin, clavulanic acid, ciprofloxaxin, cefuroxime, gentamicin, piperacillin-tazobactam, and trimethoprim. The first six of these were directly inferred from the previously published analysis. Trimethoprim was instead inferred from the *sul* and *dfrA* alleles identified by Roary; data from the Cambridge University Hospitals collection ^16^ was used to train a model constructed with the randomForest R library (https://cran.r-project.org/web/packages/randomForest/) which had 93% accuracy when applied back to the training dataset. This was used to infer resistance phenotypes for the BSAC collection.

Analysis used the heterogeneous multilocus NFDS model described previously ^17^, modified to treat a vaccine cost, *v*, as a fitness advantage, *r*. All individuals, *i*, of the sequence clusters corresponding to ST131 and ST69 were assigned the same fitness advantage, *r*_*i*_ = *r*; *r*_*i*_ = 0 for all other *i*. Hence the function defining the number of progeny, *X*_*i,t*_, produced by *i* at time *t* was:

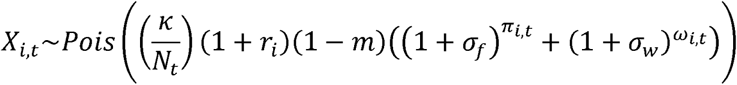

In this formula, density-dependent competition is parameterised by the carrying capacity κ, set at 50,000 to represent a large population that is still computationally feasible, and the total number of cells in the simulated population at *t, N*_*t*_. The strength of NFDS was determined by the parameters *p*_*f*_, σ_*f*_ and σ_*w*_. As previously, the accessory loci and resistance phenotypes were ordered according to the statistic Δ_*l*_:

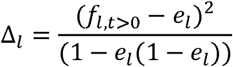

Where *f*_*l,t>0*_ is the mean post-2001 locus frequency. If the *L* loci and phenotypes considered to be under NFDS were ordered by ascending values of Δ*l*, then *l*_*f*_ was the highest ranking locus meeting the criterion 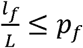. This determined the strength of NFDS acting on each locus, and therefore the reproductive fitness of individual *i*, based on which loci were encoded in its genome, as represented by the binary variable *g*_*i,l*_, and the deviation of their simulated locus frequency at time *t, f*_*l,t*_, from their corresponding equilibrium frequencies:

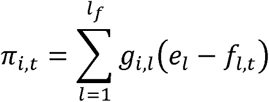

And:

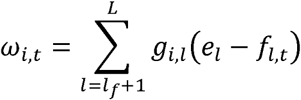

These summed deviations served as the exponents for the NFDS terms of the reproductive fitness, with π_*i,t*_ and σ_*f*_ corresponding to those loci under stronger NFDS, and ω_*i,t*_ and σ_*w*_ corresponding to those loci under weaker NFDS.

The simulations were initialised with a random selection of κ genotypes from the genomic data, which were biased such that those isolates observed in 2001 were represented at one thousand fold greater frequency than genotypes collected in later years. This was necessary to ‘seed’ the initial population with ST131 and ST69, to facilitate their expansion in a realistic manner in subsequent years. The parameter *m* represented the rate at which all isolates entered the population through migration; this was biased to import all sequence clusters at the same rate, to avoid any fits in which high rates of migration would artefactually replicate the population observed in the later years of the collection ^17^.

### Model fitting to genomic data

As in Corander et al. ^17^ the simulation model was fitted through Approximate Bayesian Computation (ABC) using the BOLFI algorithm, which has been shown to accelerate ABC inference 1000-10000 times without loss of accuracy ^30^. The prior constraints placed on the parameter values were as follows: the lower bound on all parameters was set to 0.0009 and the upper bounds were *r*_*i*_ – 0.99, *m* – 0.2, *p*_*f*_ – 0.99, σ_*f*_ – 0.03, σ_*w*_. – 0.005. We used 500 iterations of the BOLFI algorithm to minimise the Jensen-Shannon divergence of the sequence cluster frequencies in the genomic data and in the simulations, as ascertained through randomly sampling discrete sets of isolates in accordance with the size and timings of the genomes selected for sequencing from the original collection. Convergence of BOLFI was monitored each 100 iterations and the approximate likelihood estimate was assessed to have been stabilized by the end of the 500 iterations ^30^. The 95% posterior credible intervals for the parameters were obtained using three generations of sequential Monte Carlo sampling with the same default settings as used in Corander et al ^17^. The neutral model was fitted by fixing *p*_*f*_, σ_*f*_, and σ_*w*_ at zero and estimating *r* and *m* through 500 iterations of the BOLFI algorithm, followed by sequential Monte Carlo sampling, as with the full model.

## Results

### NFDS on accessory loci can explain ExPEC population dynamics

Previous work on this population suggested it was subject to balancing selection based on the persistent diversity of strains, and stable prevalence of resistance phenotypes, despite the invasion of genotypes ST69 and ST131, the latter of which has an MDR phenotype ^16^. It is possible this could represent strains being adapted to distinct niches through unique gene content. However, using the previous analysis of gene content with Roary, the 18 strains with at least ten representatives in the population had a mean of only 16.7 private genes (range: 1-49), defined as those loci present at >95% in one strain, and <5% in all others. This is consistent with strains being defined by a characteristic combination of common accessory loci, rather than distinctive sequence ^14,31^.

Such distribution of gene content is similar to that observed in *S. pneumoniae*, in which NFDS acting on variable phenotypes encoded by genomic islands was suggested to shape the population ^17^. The Roary analysis identified 6,824 intermediate-frequency genes, present in between 5% and 95% of the overall population. Comparisons between the pre-ST131 2001 samples, and subsequent data from up to 2011, found strong, linear correlations between the prevalence of their intermediate-frequency genes (Fig 1A, Fig S2). This is consistent with these loci existing at ‘equilibrium’ frequencies, determined by their costs and frequency-dependent benefits. Furthermore, these correlations with the first sample, in 2001, did not successively weaken year-on-year, as might be expected with neutral drift (Fig 1B). Instead, deviation from the first sample increased until 2008, as the sequence clusters (SCs) primarily associated with ST131 and ST69 became more prevalent (Fig 1C). The rise of ST131 was primarily driven by a dramatic rise in the prevalence of MDR clade C isolates, with clade B persisting at a lower, but stable, level. This was followed by a reversion back towards the equilibrium gene frequencies up to 2010, which does not correspond to major changes in the frequency of either ST131 or ST69, suggesting a reconfiguration of other lineages in the population.

**Figure 1:**
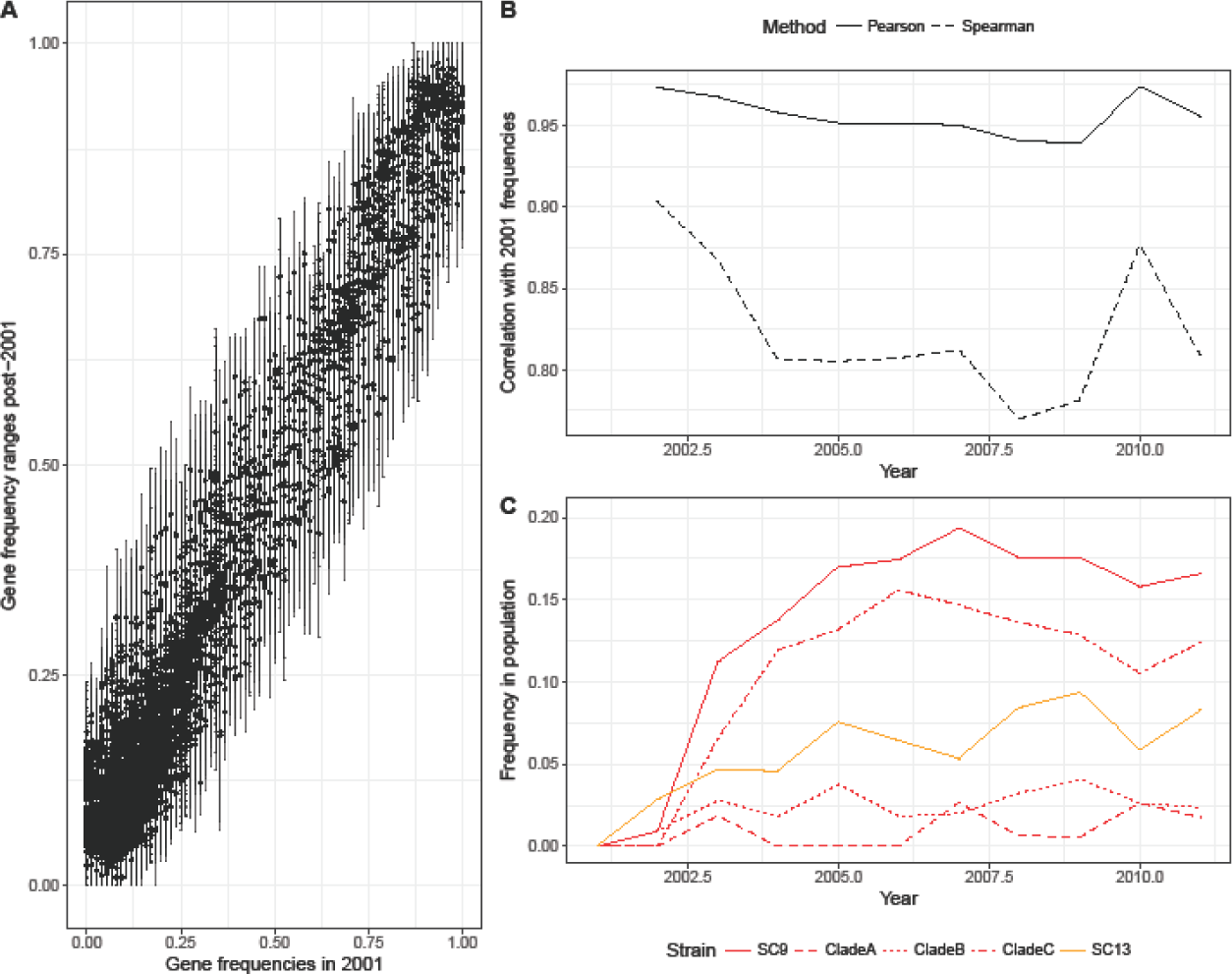
Summarising the population dynamics of the British Society for Antimicrobial Chemotherapy extraintestinal pathogenic *E. coli* collection. These isolates were collected from bacteraemia cases around the UK between 2001 and 2011. (A) Conservation of gene frequencies. Each point corresponds to one of the 6,824 genes identified by ROARY in the BSAC collection with a mean frequency between 0.05 and 0.95 across all years. The horizontal axis position indicates the starting frequency in 2001, and the vertical axis indicates the mean frequency over all years, with the error bars indicating the full range observed across annual samples. (B) Correlation of gene frequencies with those observed in 2001. This shows the changing correlation of gene frequencies, calculated by both the Pearson and Spearman methods, in each year relative to those observed in 2001. Both measures indicate a divergence in gene frequencies as ST69 and ST131 emerge, until 2010, at which point there is a reversion to the frequencies seen in the original population. (C) Emergence of ST69, in orange, and ST131, in red. The frequencies of the subclades of ST131 are shown by the red dashed lines.

In order to obtain a population-wide view of these dynamics, the previously-described multilocus NFDS model was applied to this dataset to test whether these strain dynamics were consistent with selection at the accessory locus level. The model was initialised with the 2001 population, which was seeded with genotypes observed in later years at a low level, representing the possibility they were present in the population but unsampled. Subsequent simulation with a Wright-Fisher framework included these post-2001 genotypes migrating into the population at a rate *m*, while the hierBAPS clusters corresponding to ST131 and ST69 expanded at a rate determined by their increased reproductive fitness relative to the rest of the population, *r*. The equilibrium frequencies of 7,211 intermediate frequency loci, corresponding to genes identified by Roary that were between 5% and 95% in the 2001 sample plus ten antibiotic resistance phenotypes, were assumed to be those observed in 2001 sample of genomes. These were then simulated as evolving under

NFDS; a fraction *p*_*f*_ evolved under strong NFDS, determined by the parameter σ_*f*_, while the rest evolved under weak NFDS, according to parameter σ_*w*_ (see Methods). Fitting this model using BOLFI estimated the parameters listed in Table S2, which identified significant evidence for NFDS (σ_*f*_ and *p*_*f*_ greater than zero), providing a gene-level mechanistic basis for NFDS underlying the previous strain-level observations of Kallonen *et al* ^16^.

These simulations successfully reproduced several aspects of the observed data (Fig 2, Fig S3). Both ST131 and ST69 rapidly spread through the population, before stabilising at an equilibrium frequency. This does not occur at the expense of the established, common clones, such as ST73 and ST95. Instead, in accordance with the genomic data, the displaced sequence clusters include ST10, ST14, ST144 and ST405. These patterns are qualitatively distinct from an equivalent neutral model fit (Fig 2C). Without NFDS, both ST131 and ST69 are predicted to exponentially increase in frequency, with all other strains decreasing at accelerating rates, proportionate to their original prevalence. The greater invasion rate of ST131, relative to ST69, is an artefact of its higher prevalence in the overall dataset meaning it is seeded at a higher level, rather than a true ecological difference. Although NFDS constrains the invasion of new strains in these simulations, the multidrug-resistant clade C of ST131 is still able to reach high prevalence, even when such selection is active. This may be at least partially attributable to some members of this recently-emerged clade C having considerably diversified in their genome content, as indicated by pairwise comparisons of gene content, which show clade C isolates were similar to those between random representatives selected from the same sequence cluster (Fig S4). This might enable the clade to avoid the limitations of any loci that NFDS would suppress to low frequencies. Hence the underlying genotype of ST131 appears to represent a highly-fit genotype that has subsequently diversified into both antibiotic-sensitive (clade B) and resistant (clade C) forms, expressing one of multiple capsules ^35^. Therefore a comprehensive genomic dataset encompassing all known ST131 genome sequences was created to understand the unique characteristics of the ST131 lineage, with particular focus on the successful clades B and C.

**Figure 2:**
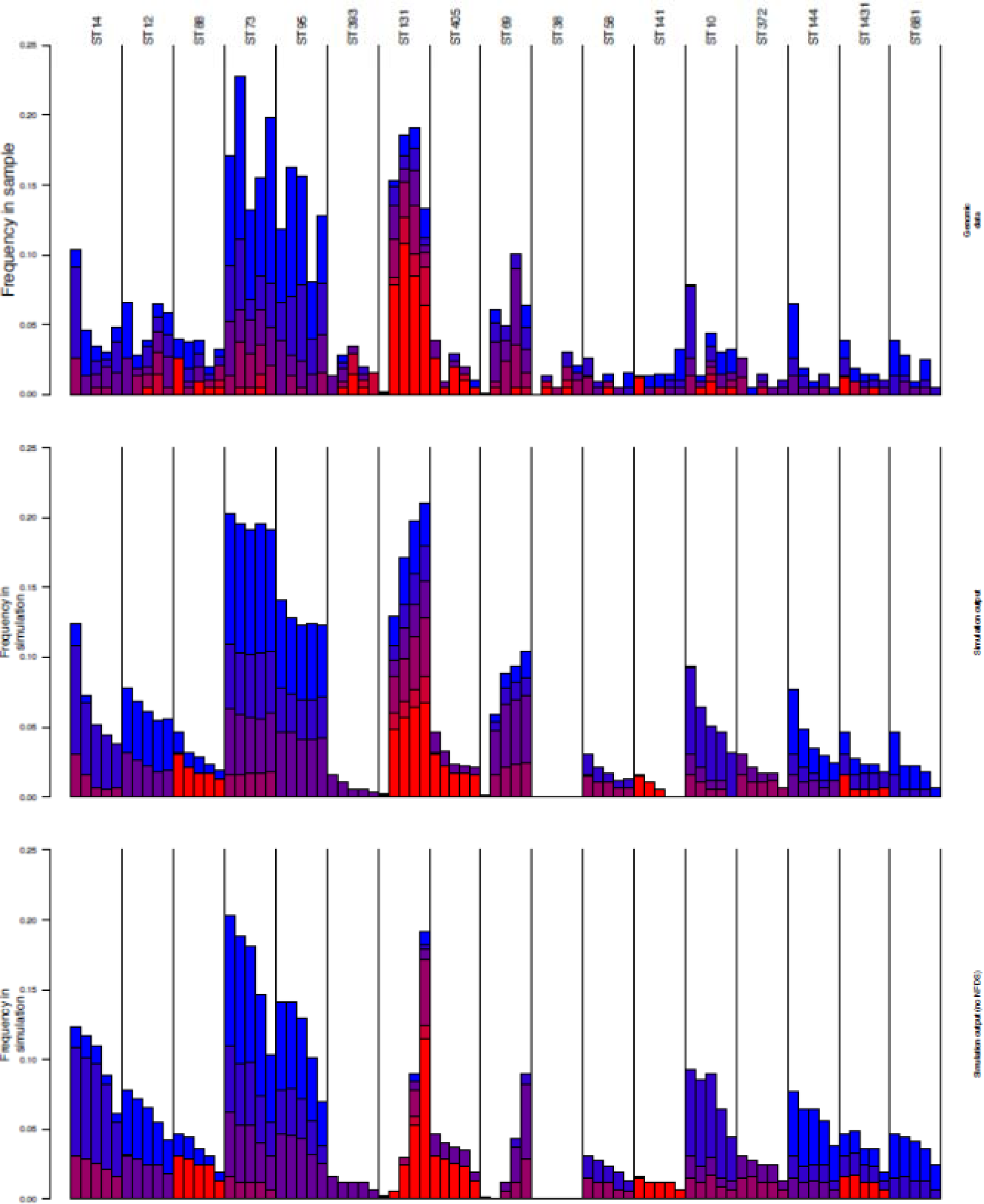
Simulations of changes in the BSAC extra-intestinal pathogenic *E. coli* population evolving under multilocus NFDS. Panel A shows the genomic data, and panel B shows the median frequencies observed from 100 simulations run with the best-matching parameter set identified by fitting the model with BOLFI. This corresponded to σ_*f*_ = 0.029, *r* = 0.179, *m* = 0.001, *p*_*f*_ = 0.425 and σ_*w*_ = 0.0048. Each column corresponds to a sequence cluster identified by hierBAPS (see Methods), and is annotated with the predominant sequence type with which it is associated. Each bar indicates the frequency of the sequence cluster in consecutive time periods, from left to right. The bars are coloured according to the number of antibiotic resistance phenotypes associated with the isolates within the sequence cluster at different timepoints. Panel C shows the equivalent best fit in the absence of NFDS. Only sequence clusters reaching a frequency of at least 2.5% at one timepoint in the genomic sample are shown; the full results of the simulation, including measures of between-simulation variation, are shown in Fig S3.

### Core and accessory genomic structure of the ST131 population

A maximum likelihood phylogeny generated from an alignment of concatenated core CDS from all 862 genomes confirmed the earlier consensus three clade structure of the lineage (Fig 3a), and in agreement with previous studies, there was no strong phylogeographic signal or host source clustering evident in the phylogeny (https://microreact.org/project/BJKoeBt2b). To confirm that the collation of the 862 genomes was consistent with previous descriptions of the accessory genome distribution in ST131, isolate relatedness based on shared accessory gene content was visualized as a two-dimensional projection using PANINI (Fig 3b) ^24^. Clades A and B largely resided in dense clusters at the periphery of the projection. In contrast, clade C isolates were more diffuse, overlapping with some clade B isolates, forming a cloud with discernible sub-structuring into distinct groups. This concurs with the previous analysis of the gene content of clade C, and the previous finding of multiple accessory genome sub-clusters in clade C ^14^.

**Figure 3:**
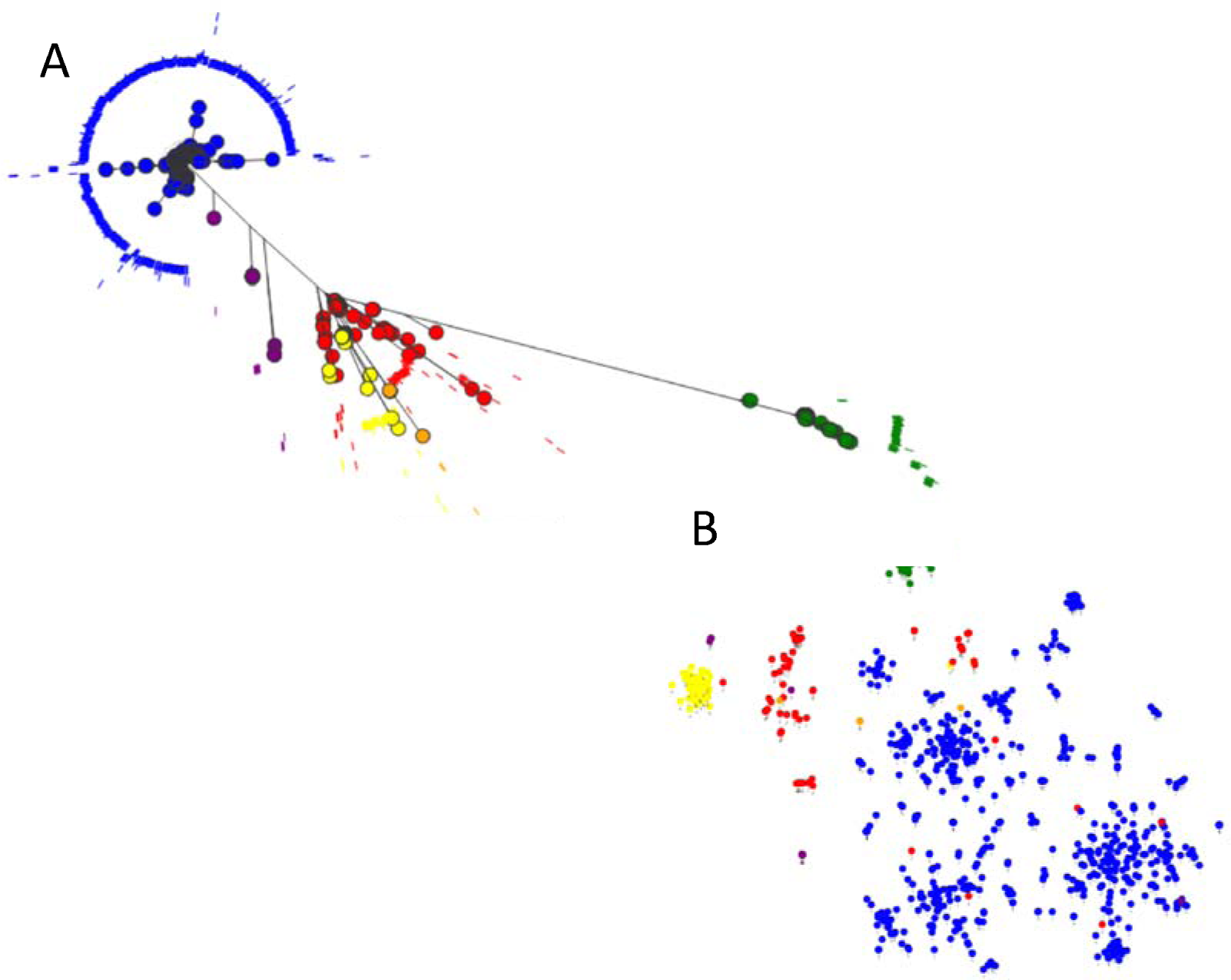
(A) Maximum likelihood phylogeny of 862 *E. coli* ST131 strains. The phylogeny was inferred using RAxML with a GTR GAMMA model of substitution, on an alignment of concatenated core CDS as determined by Roary. (B) PANINI plot of the accessory genome content of all 862 strains based on a tSNE plot. The plot is a diagrammatical representation of the relatedness of each strain based on the presence/absence of accessory genes, and is presented as a two dimensional representation. The taxa are colour coded by BAPS grouping (Table S1) and show clade A (Green, BAPS-3), clade B (red, yellow and purple – BAPS 2, 4, and 5) and clade C (blue, BAPS-1).

### Low frequency accessory genes suggest differential ecology of clade A and clade B/C *E. coli* ST131

Given that the vast majority of accessory genes occur at very low frequency, we sought to determine if these represented mobile genetic elements circulating transiently in the population. We functionally categorised genes occurring in less than 20% of the overall ST131 sample (based on the distribution of the gene frequencies in Fig S1) that were confined to a single clade. In both clade A and clade B/C (Dataset S1-S3) the overwhelming majority of low frequency accessory genes encode hypothetical proteins (64.4% clade A, 58% clade B/C). Excluding the hypothetical proteins from the analysis showed unexpected bias in functional gene categories differentially observed in the lineages (Fig 4). The most common gene types were functional phage, plasmid and other mobile genetic element (MGE) genes, with more private phage genes present in clade B/C than in clade A. Conversely, there were more private plasmid genes in clade A than clade B/C, despite the presence of a diverse number of MDR plasmids within clade C.^14^ Together this suggests that clade A strains of *E. coli* ST131 and clade B/C strains of *E. coli* ST131 are exposed to different plasmid and phage pools, an observation which is most parsimoniously explained by them having different ecological habitats.

**Figure 4:**
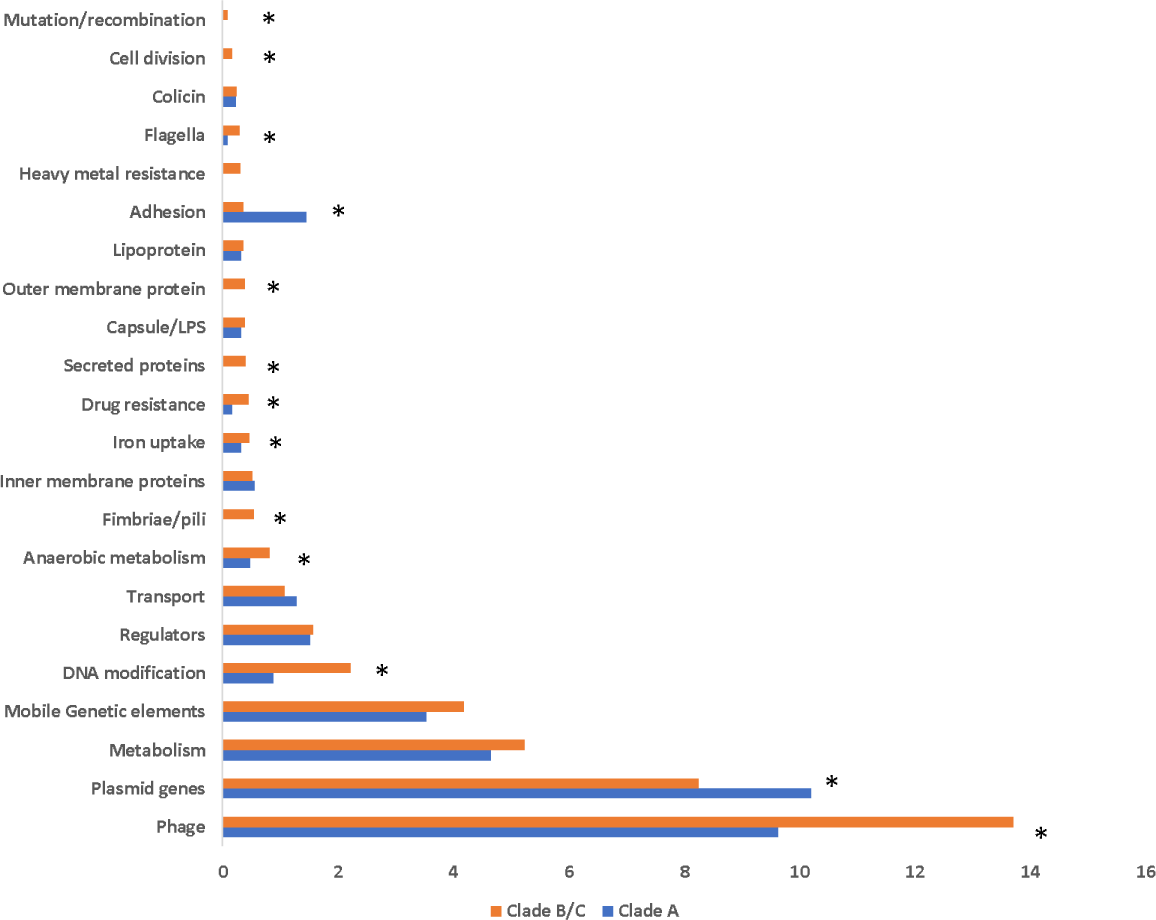
Bar chart depicting functional classes of accessory genes differentially present in clade A (blue bars) and clade B/C (orange bars) *E. coli* ST131. Functional classes are based on GO classes as described in methods. Bars marked with * indicate where a significant difference exists between clade A and clade C as determined by t-test.

### Clade-specific and intermediate frequency genes in the population

To identify which aspects of the accessory genome differed between the clades of ST131, the distributions of the 32,631 sets of orthologous genes identified by Roary were analysed (Dataset S3). Characterising the full set of loci present at intermediate frequencies was not feasible, as even focussing on the 3,354 present at between 5% and 95% frequency found the majority of these were present at a frequency below 20% (Fig S1). Therefore, the search was refined to clade specific genes, occurring at a frequency > 95% in one clade but at <5% in the other two clades (Dataset S1).

Clade A contained the highest number of loci exclusive to a lineage (54) despite constituting the least sampled clade. Clade B had only 2 exclusive loci and clade C had 18. When clades B and C were combined against clade A, there were 60 loci exclusively present in the B/C combination. The majority of clade A private genes encode hypothetical proteins whilst those private to clade C encode DNA modification proteins and metabolic functions. The genes private to clade B/C combined also encode hypothetical proteins and metabolic functions, notably five dehydrogenase enzymes involved in anaerobic metabolism labelled *yihV, garR_3, fadJ, fdhD,* and *gnd* in our dataset (Dataset S2). Blast analysis against the NCBI non-redundant database suggested that the dehydrogenase enzyme gene annotated as *pdxA* in our Roary dataset was confined to clade C ST131 strains. These dehydrogenase enzyme genes were found to be present across phylogroup B2 *E. coli* strains (of which ST131 is a member) through BLASTN searches of the NCBI non-redundant database. Therefore these loci are not unique to clade C ST131, and were either acquired by an ancestral clade B/C strain, or have been lost by clade A.

### High diversity in core anaerobic metabolism genes unique to clade B/C

Analysis of accessory loci private to clade B/C (present in >95% of that population) identified two separate loci encoding 3-hydroxyisobutyrate dehydrogenase enzymes, and loci encoding 3-hydroxyacyl-CoA dehydrogenase, 6-phosphogluconate dehydrogenase, and formate dehydrogenase. Analysis of clade B/C loci circulating at low frequency of <20% also identified a significant over-representation of genes encoding dehydrogenase enzymes involved in anaerobic metabolism (a total of 64 loci), including seven variants of formate dehydrogenase. There were also seven variants of the *eutA* gene found in the ethanolamine utilisation pathway (the *eut* operon) and a distinct version the *cobW* gene which encodes the sensor kinase for activation of the cobalamin biosynthesis operon. Closer investigation of the sequences of these loci suggested that these were not genes private to clade B/C per se, but rather represented multiple unique alleles of genes that are core to the ST131 population which differ at nucleotide sequence level by more than 5%. This implies a unique selection pressure is acting on these core genes in clade B/C compared to clade A.

Further scrutiny of low frequency loci in clade B/C also identified alternative alleles of a large number of well characterised extra-intestinal pathogenic *E. coli* virulence-associated genes, including: antigen 43 (7 alternative alleles); heavy metal resistance such as arsenic (5 loci), copper (4 loci), and mercury (5 loci); capsule biosynthesis (20 loci); cell division and septation (14 loci); antibiotic resistance to chloramphenicol (3 loci), macrolides (2 loci), rifampicin (1 locus), and MDR efflux pumps (21 loci); iron acquisition (39 loci); curli and type I fimbriae and P pili (42 loci); lateral and classical flagella (26 loci); and LPS synthesis (9 loci). These loci represent alternative alleles of genes found widely across the *E. coli* phylogeny indicating there are multiple allelic variants of important genes that are confined to clade B/C of the *E. coli* ST131 lineage.

We sought to determine the distribution of this allelic diversity across the *E. coli* ST131 phylogeny by annotating the tips of the phylogenetic tree with the presence/absence of each of the anaerobic metabolism (Figure 5), and capsule, cell division, MDR efflux, iron acquisition, pili, and flagella divergent loci (Figure 6). Our analysis shows that each alternative allele occurs at very low frequency but that alleles are randomly distributed throughout the phylogeny of the C clade, and are exclusive to clade C. Given that these alleles differ from the normal conserved versions of genes by >5% at nucleotide level, it is implausible that these alleles would be arising repeatedly and independently via mutation. Instead, the most parsimonious explanation is that the minor frequency alternative alleles are being distributed through the population via recombination. This conclusion is supported by the fact that every one of the allele variants identified in our analysis has 100% nucleotide identity matches with genes present in other *E. coli* in the NCBI non-redundant database.

**Figure 5:**
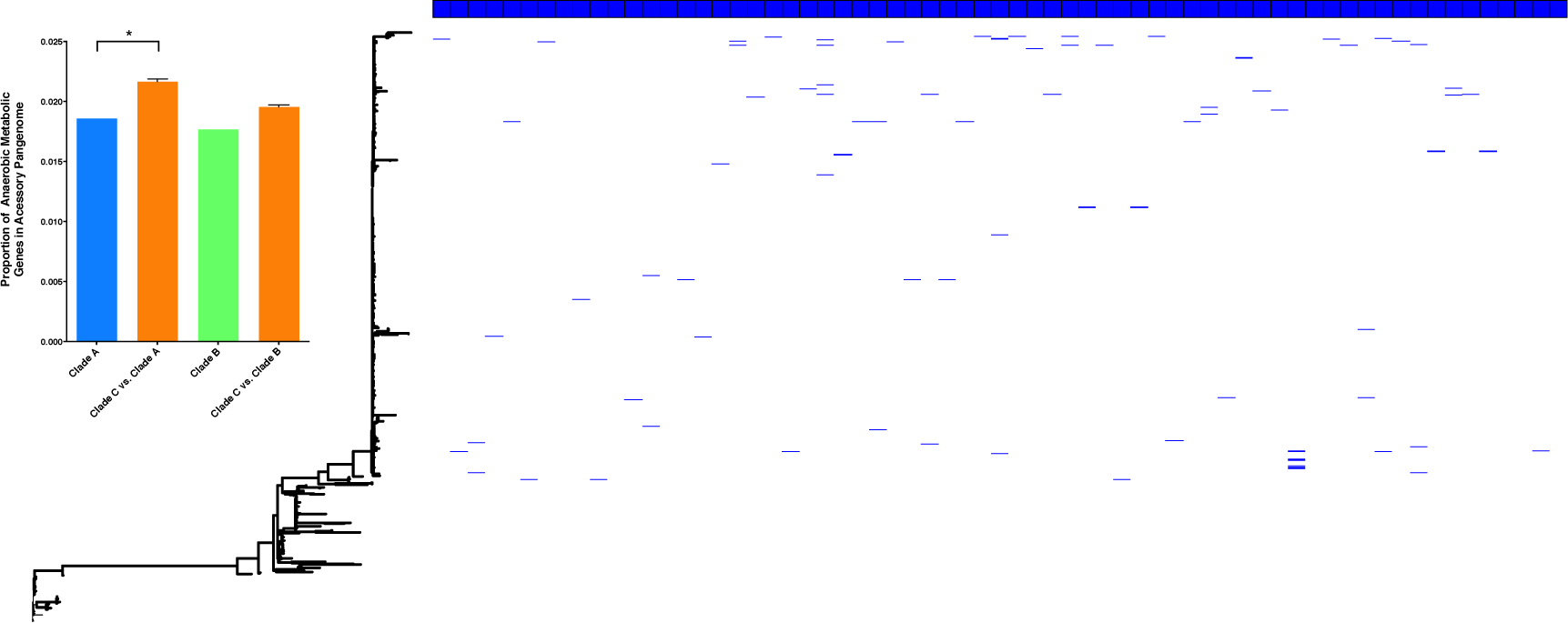
Annotation of a maximum likelihood phylogeny of *E. coli* ST131, based on concatenated core CDS, with the presence of alternative alleles of 64 loci involved in anaerobic metabolism. Each blue box along the top of the tree annotation represents an individual anaerobic metabolism gene, and its presence in the ST131 population is indicated by a blue line. The inset is a bar chart displaying the proportion of the accessory pangenome that is occupied by genes involved in anaerobic metabolism for ST131 Clade A (light blue), Clade B (light green), subsampled Clade C vs. Clade A (orange) and subsampled Clade C vs. Clade B (orange). P = 0.042 for Clade C vs. Clade A and P = 0.086 for Clade C vs. Clade B. Error bars represent standard error of the mean. Significance was determined using the median value p-value from Chi squared tests performed on random subsamples of the C clade.

**Figure 6:**
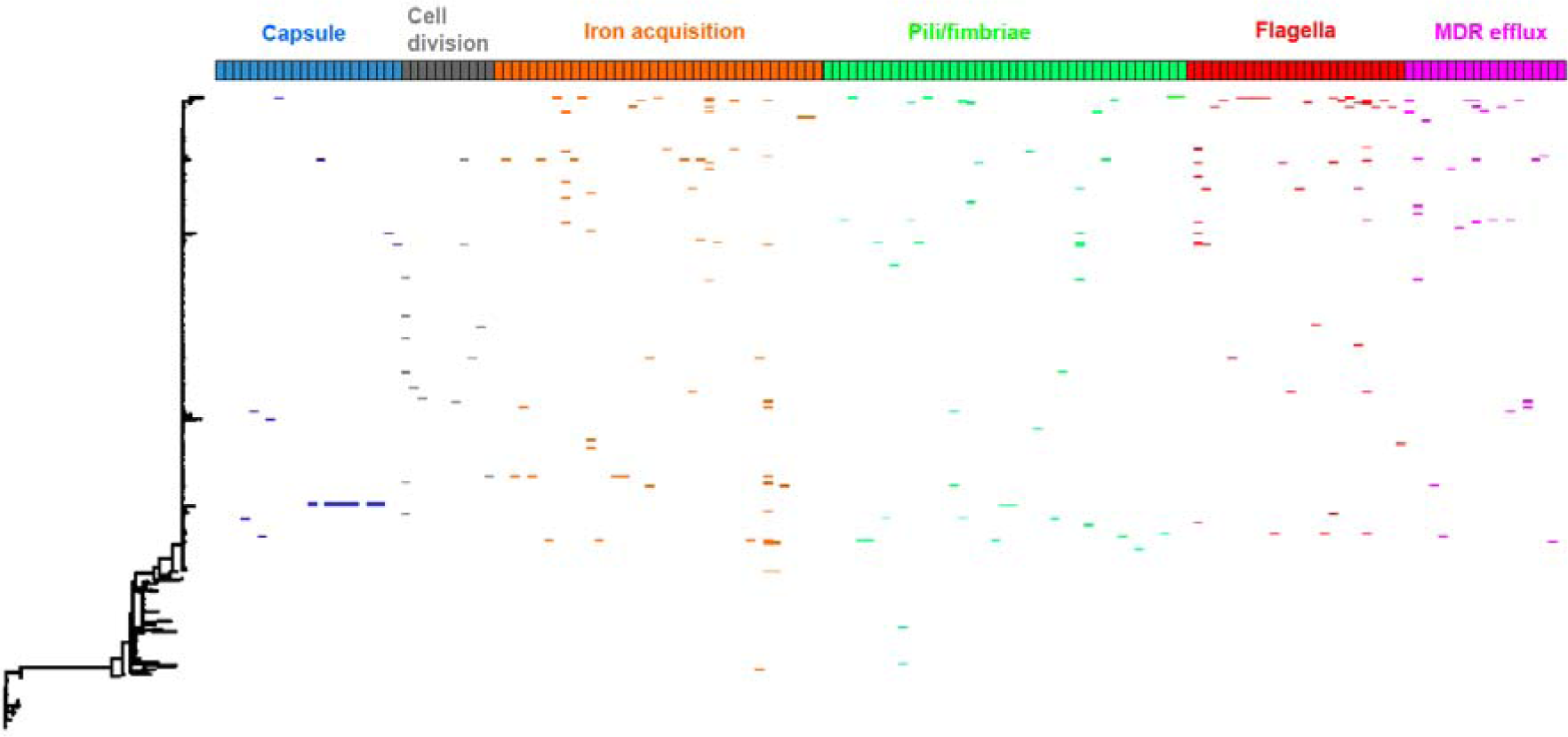
Annotation of a maximum likelihood phylogeny of *E. coli* ST131, based on concatenated core CDS, with the presence of alternative alleles of loci involved in capsule production (blue boxes), cell division (grey boxes), iron acquisition (orange boxes), pili/fimbriae production (green boxes), flagella (red boxes), and MDR efflux pumps (pink boxes). Each box represents an individual gene, and its presence in the ST131 population is indicated by an appropriately coloured line.

Given that our data set is biased towards clade C genomes, we performed comparative analyses of the frequency with which allelic diversity occurs in anaerobic metabolism genes. We randomly subsampled clade C 100 times and compared an equal number of clade A, B, and C genomes for allelic diversity. Our data shows that even when randomly subsampling clade C, the levels of diversity observed in anaerobic metabolism genes is significantly higher than in clade A, providing evidence that the accumulation of sequence diversity is specific to the MDR clade C (Figure 5).

Finally, we sought to exclude the possibility that the presence of these allelic variants was skewed by some form of geographically localised expansion of variants. To do this we compared the relative frequency of all accessory genes, highlighting the allele variants in anaerobic metabolism, capsule, cell division, MDR efflux, iron acquisition, fimbriae, and flagella present in UK versus non-UK isolate genomes (Figure S5). Our data showed a strong linear relationship between the frequency of genes in the two populations, indicating that the data was not biased by expansion of alleles in a given geographical location, and that this accumulated diversity was equally as likely to happen in any given strain independent of its geographical origin.

### Allelic diversity of anaerobic metabolism genes in Clade C ST131 is not observed in other dominant ExPEC lineages

The possibility exists that the above observations made for clade C of *E. coli* ST131 simply reflect the general evolutionary path of a successful extra-intestinal pathogen. To test this we performed an identical analysis on the pangenome of 261 ST73 isolates and of 160 ST95 isolates from the UK BSAC population survey ^16^. *E. coli* ST73 and ST95 represent two of the most dominant lineages associated with clinical extra-intestinal disease alongside ST131 ^5,16^, but are predominantly non-MDR lineages and rarely associated with MDR plasmids ^16^. As with our inter-clade comparisons, we randomly subsampled clade C ST131 100 times to allow equal numbers of genomes per lineage to be compared. Our analysis showed a similar ratio of plasmid, phage and hypothetical proteins in the accessory genome as in ST131 (Fig 7). ST73 and ST95 displayed similar ratios of alternative alleles in P and Type 1 fimbriae, cell division and septation genes, and multiple iron acquisition genes as observed in ST131. However, enrichment in allelic variation in anaerobic metabolism genes was significantly higher in any given subsampled set of clade C ST131 genomes compared to both lineages. This supports the hypothesis that the observation of increased diversity accumulating in anaerobic metabolism genes is not a more general extra-intestinal pathogenic *E. coli* trait but is particularly enriched in the ST131 lineage.

**Figure 7:**
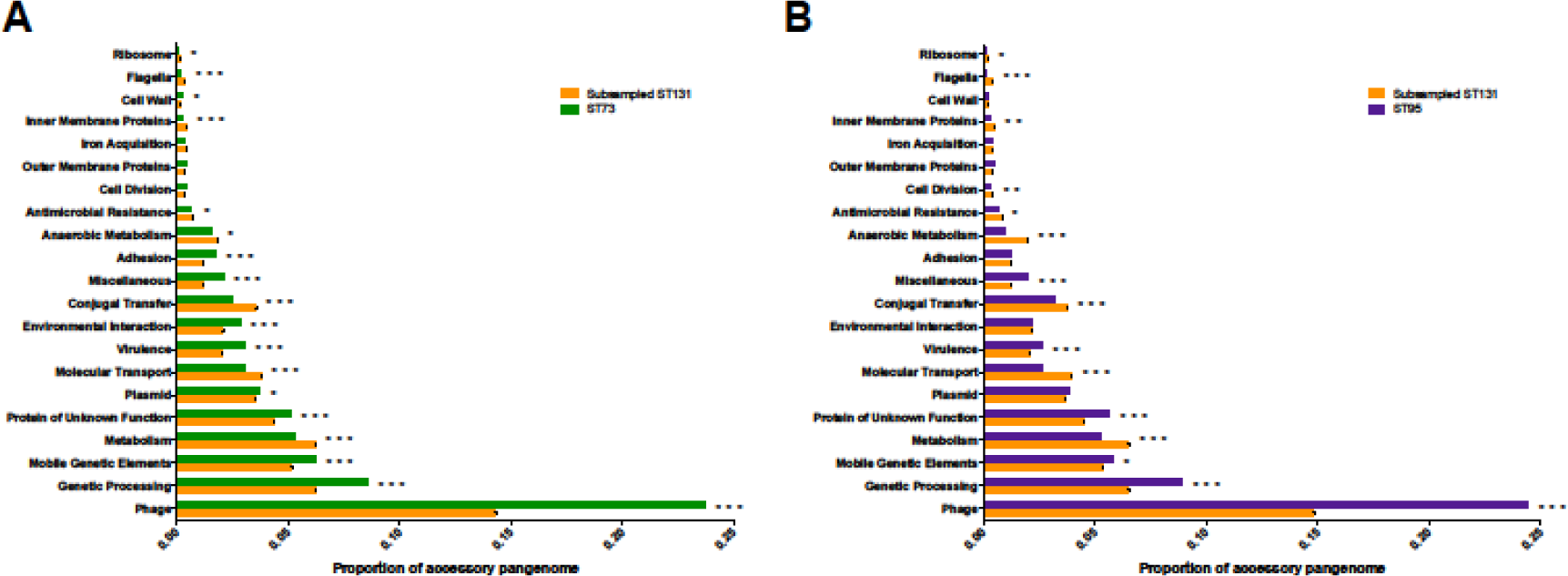
Bar charts depicting the composition of the accessory genome of ST73 (green) and ST95 (purple) compared to a repetitively sampled Clade C ST131 (orange). The proportion of the accessory genome is plotted against manually assigned functional categories. Hypothetical proteins are responsible for the majority of the accessory pan genome and are omitted from the graphs. Error bars are standard error of the mean. Iterative Chi squared tests were performed to assess significance, as described in methods, p<0.05 (*), p<0.01 (**) and p<0.001 (***).

The accumulation of nucleotide diversity in a given set of loci can often be interpreted as a signature of some form of selection occurring on those genes. However the low levels of frequency of any given allele across clade C strains contradicts a hypothesis for positive selection, where one would expect successful or beneficial alleles to sweep to a high frequency or fixation. Indeed comparison of the sequences of each of the 64 anaerobic metabolism loci in which diversity was observed identified just three loci which showed signatures of positive selection as indicated by a Tajima’s D score above two.

However, these results can be reconciled with a lineage evolving under NFDS. Different resource use strategies can facilitate co-existence between competing strains, such those co-colonising a host, resulting in frequency-dependent selection ^32,33^. This would explain the sustained intermediate frequencies of genes encoding dehydrogenases over multiple years (Fig S6). Hence this diversification of metabolic loci could represent the adaptive radiation of particular traits within a successful genetic background, able to efficiently compete with the resident *E. coli* population through a diverse panel of metabolic capacities suited to exploiting resources under anaerobic conditions.

## Discussion

The evolutionary events that led to the emergence of *E. coli* ST131 have been an intense focus of research, with consensus opinion suggesting that, following acquisition of key ExPEC virulence factors, acquisition of fluoroquinolone resistance in the 1980’s by the clade C sub-lineage of ST131 was a key event in that emergence ^11,12^. However, a recent nationwide UK population survey rejected this hypothesis and suggested that success of the major ExPEC clones is not dictated by resistance traits ^16^. Here, we identify the conserved frequencies of accessory genes in the *E. coli* population which strongly suggest this species’ population structure and dynamics are shaped by NFDS acting on genomic islands. Such multilocus NFDS is able to account for how an otherwise stable population was disrupted by the invasion of ST131 and ST69, displacing some lineages while leaving other, largely antibiotic-susceptible, genotypes at almost untouched prevalence.

Previous work has suggested that clade C strains of *E. coli* ST131 undergo reduced levels of detectable core genome recombination compared to other phylogroup B2 *E. coli* ^36^ or ST131 clade A strains ^14^. We have previously postulated that this may be a result of ecological separation between clade C strains and other common ExPEC^14,36^. Our analysis of nearly 900 genomes has allowed us to interrogate accessory gene movement to a far greater resolution than previously possible. From the analysis of the accessory genome we identified thousands of plasmid, phage and other mobile genetic element genes which are private to clade A and the combined clade B/C, respectively. Such an observation is a classic signature of ecological separation of the two populations ^37,38^, particularly given that the genetic distance between clade A and clade B/C is much smaller than it is to other lineages and species from which the circulating genes are also found in the NCBI non-redundant database.

Our analysis also identified a significantly increased level of sequence diversity in genes involved in key host colonisation processes in clade C. This diversity was uncovered through our pan-genome analysis as allelic variants of core genes. Primary amongst these is a large number of genes involved in anaerobic metabolism, including seven allelic variants of the formate dehydrogenase gene, as well as allelic variants of genes involved in ethanolamine utilisation and cobalamin biosynthesis. The pivotal role of ethanolamine production and cobalamin biosynthesis in the ability of Gram negative pathogens to outcompete bacteria in the human intestine is well documented ^39,40^, and this phenomenon only occurs when supported by an increased ability to perform anaerobic respiration in the presence of inflammation ^39^. It has been shown that MDR *E. coli* ST131 is able to colonise the gastro-intestinal tract of humans for months or years in the absence of antibiotic selection ^41,42^, and that this colonisation results in a displacement of the *E. coli* colonising the host prior to exposure to the MDR strain ^41^.

Whilst this diversity in anaerobic metabolism genes was unique to clade C ST131, the allelic variation observed in other human colonisation and virulence factors such as iron acquisition, fimbriae, and cell division was also observed in two of the other most commonly isolated lineages of *E. coli* from extra-intestinal infections, ST73 and ST95. This diversity likely reflects selection occurring on genes important for ExPEC pathogenesis. Iron acquisition is well characterised as a key virulence determinant in ExPEC, with the ability to initiate a successful UTI completely abrogated in the absence of functional iron acquisition systems ^43^. Recent experimental vaccine work exploiting siderophore production by ExPEC has shown to be highly effective in rodent models on ExPEC UTI ^44^. The importance of iron acquisition can also explain many of the MDR efflux allele variants seen in this data set, with half occurring in the *acrD* gene which has been experimentally shown to play a role in iron acquisition in *E. coli* ^45^. We identified multiple alleles of genes in the type 1 fimbriae operon and in genes in the P pilus operon which are classical virulence determinants in UTI ^46^, and multiple genes involved in capsule biosynthesis, which we have previously reported as being a hotspot for recombination in *E. coli* ST131 ^13,35^. We also identified multiple alleles of genes involved in controlling incomplete septation and filamentous growth, which is a crucial process in the formation of the filamentous intracellular bacterial communities (IBCs) which are thought to be fundamental in the ability of ExPEC to survive inside bladder epithelial cells and cause UTI ^47^. There are a small number of allelic variants in anaerobic metabolism genes also present in ST73 and ST95, possibly reflecting recent experimental studies suggesting a crucial role for the cytochrome-bd oxidase system in the ability to cause urinary tract infection ^48^. Also previous studies using saturated mutagenesis techniques and studying global transcriptional patterns during urinary tract infection of ExPEC strains have suggested a key role for dehydrogenase enzymes involved in anaerobic metabolism in the ability to cause pathology in the mammalian urinary tract ^49–51^.

Recent modelling data on why drug resistant and drug susceptible populations of bacteria co-exist highlighted that any factors which increase the duration of colonisation in a human host will also increase the selective pressure for it to evolve antibiotic resistance ^52^. Hence both the success of ST131 in invading the population, and the association of many isolates in this lineage with an MDR phenotype, would be consistent with its distinctive anaerobic metabolism loci facilitating enhanced persistence within its host, perhaps through an improved ability to outcompete resident commensal *E. coli* strains. The fact that this selection is only seen in clade C of ST131 suggests that this occurred around the time of the emergence of the lineage as a human clinical threat ^13^ alongside the development of fluoroquinolone resistance. Subsequent acquisition of MDR plasmids, and the consequent selection for an ability to offset the fitness costs of long term MDR plasmid maintenance ^14^, is likely to have occurred as a result of prolonged exposure to selective antibiotic environments during colonisation of humans. Nevertheless, neither anaerobic metabolism genes nor antibiotic resistance loci have swept to fixation in ST131, reflecting their fluctuating but stable prevalence in the broader *E. coli* population (Fig S6).

This diversification can instead be explained by NFDS, under which these genes are beneficial when rare, because they provide an advantage over co-colonising strains which will typically lack the same metabolic capacities. However, as these traits become more common as ST131 expands, representatives of this lineage will more commonly encounter one another, therefore necessitating further diversification for different clade C representatives to sustain their advantage over competitors. Similarly, the capsule locus diversification previously observed within clade C, resulting in the capsule synthesis locus corresponding to a ‘hotspot’ of recombination^35^, could result from NFDS of variable antigens ^54^, with the host immune system selecting for a diversity of capsule structures as the dominant type becomes more common following ST131’s emergence^16^.

This study presents evidence for both ecological niche separation, resulting in the formation of distinct subclades within ST131, and NFDS, resulting in the adaptive radiation of specific phenotypes within clade C as it increases in prevalence. Further studies are required to fully determine the extent to which niche separation and NFDS are either separate or linked processes. Determining whether loci subject to NFDS are also those that determine niche adaptation will be integral to this process. Understanding the processes that govern the epidemiological dynamics of dominant *E. coli* lineages, and those of similar pathogens causing bloodstream infections, is critical for addressing the public health threat of antibiotic resistance.

## Data accession

Accession numbers for the reads used in this study are listed in Table S1 with information of year and place of isolation and the results of the *in silico* PCR for clade specific SNPs.

## Acknowledgements

This publication presents independent research supported by the Health Innovation Challenge Fund (HICF-T5-342 and WT098600), a parallel funding partnership between the UK Department of Health and Wellcome. The views expressed in this publication are those of the authors and not necessarily those of the Department of Health, Public Health England or Wellcome. T.K. was funded by the Norwegian Research Council JPIAMR grant no. 144501. J.C. was funded by the ERC grant no. 742158. NJC is funded by a Sir Henry Dale Fellowship, jointly funded by the Wellcome Trust and Royal Society (Grant Number 104169/Z/14/Z). CC is funded by the Wellcome MIDAS doctoral training program at UoB.

---

**Figure S1:**
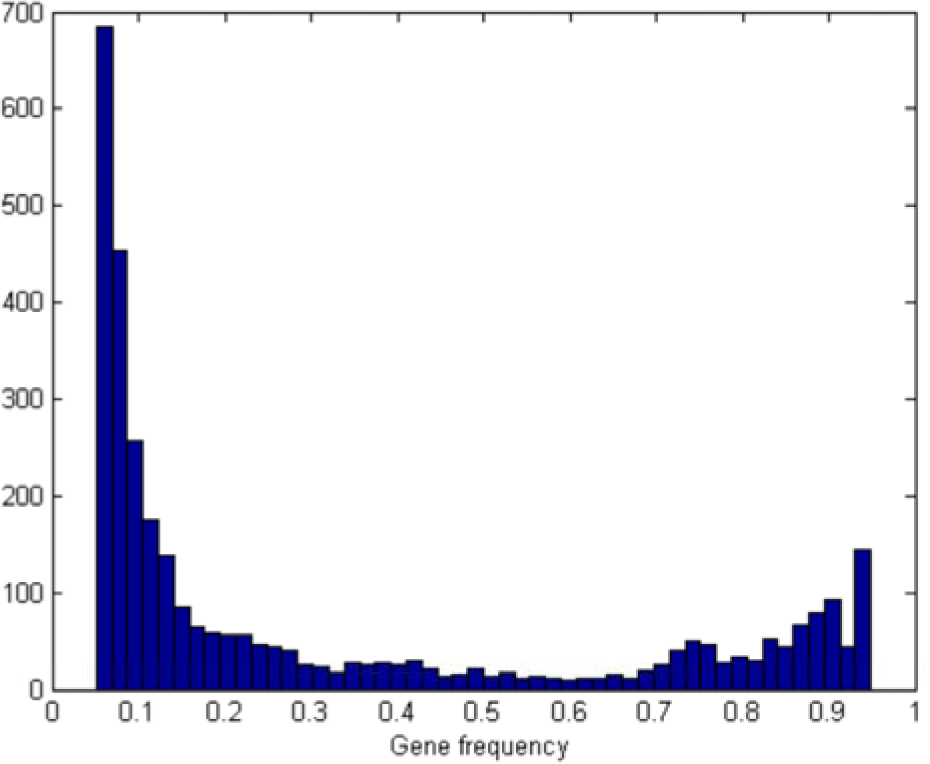
Histogram of the relative frequency of genes within the accessory genome of *E. coli* ST131. The x-axis indicates the relative frequency with which a gene appears, whilst the y-axis indicates the number of accessory genes which appear at that given frequency.

**Figure S2:**
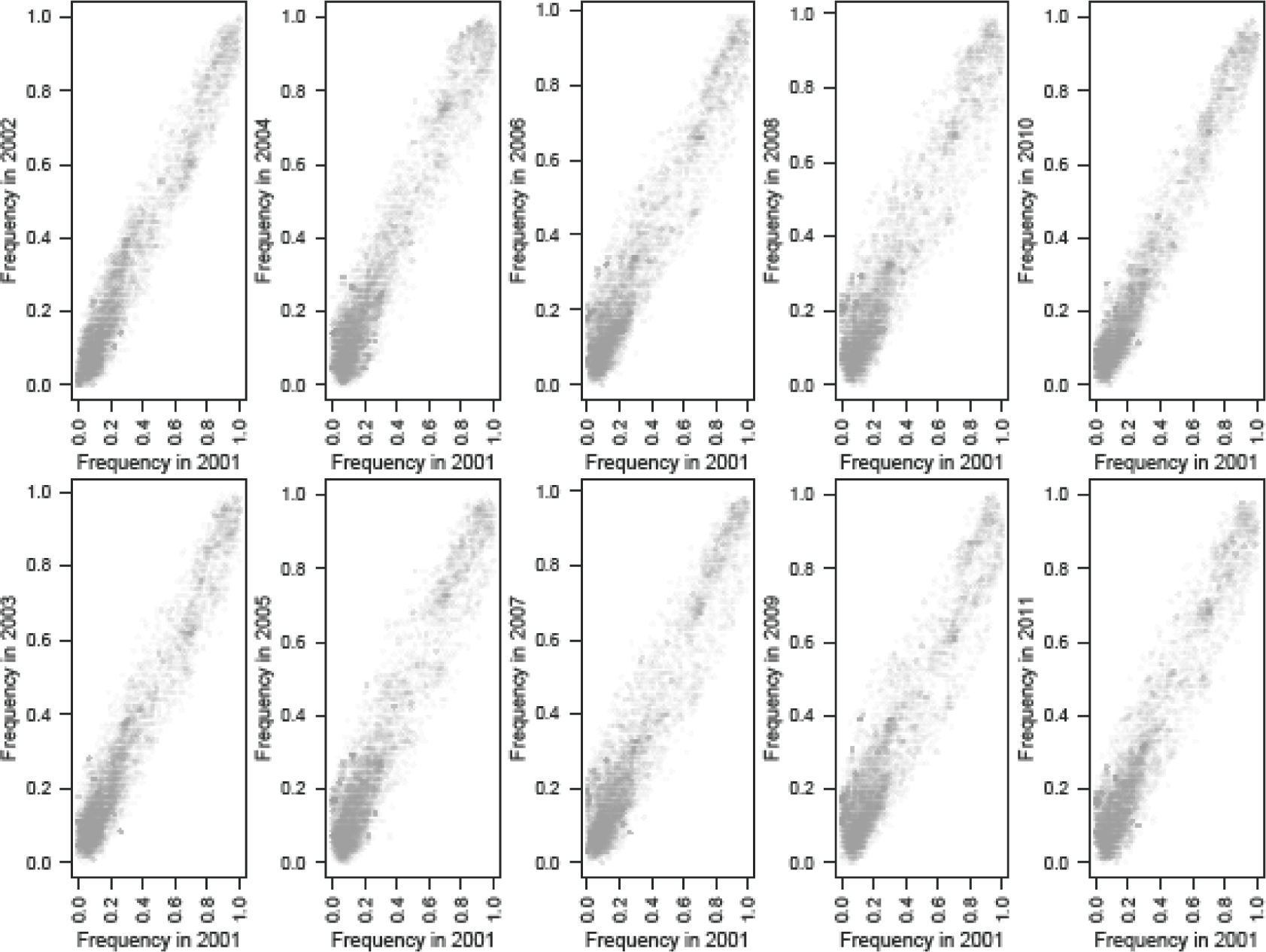
Correlations of gene frequencies in the BSAC collection over time. Each plot shows the frequencies of those genes, identified by ROARY, that were found to be present at a mean frequency between 0.05 and 0.95 across the entire collection. In each panel, the horizontal axis shows the frequency in 2001, and the vertical axis shows the frequency in a subsequent year. These graphs show how the correlation between the starting frequencies, in 2001, and later years weakened until 2008, at which point the correlation strengthen considerably in 2010 and 2011.

**Figure S3:**
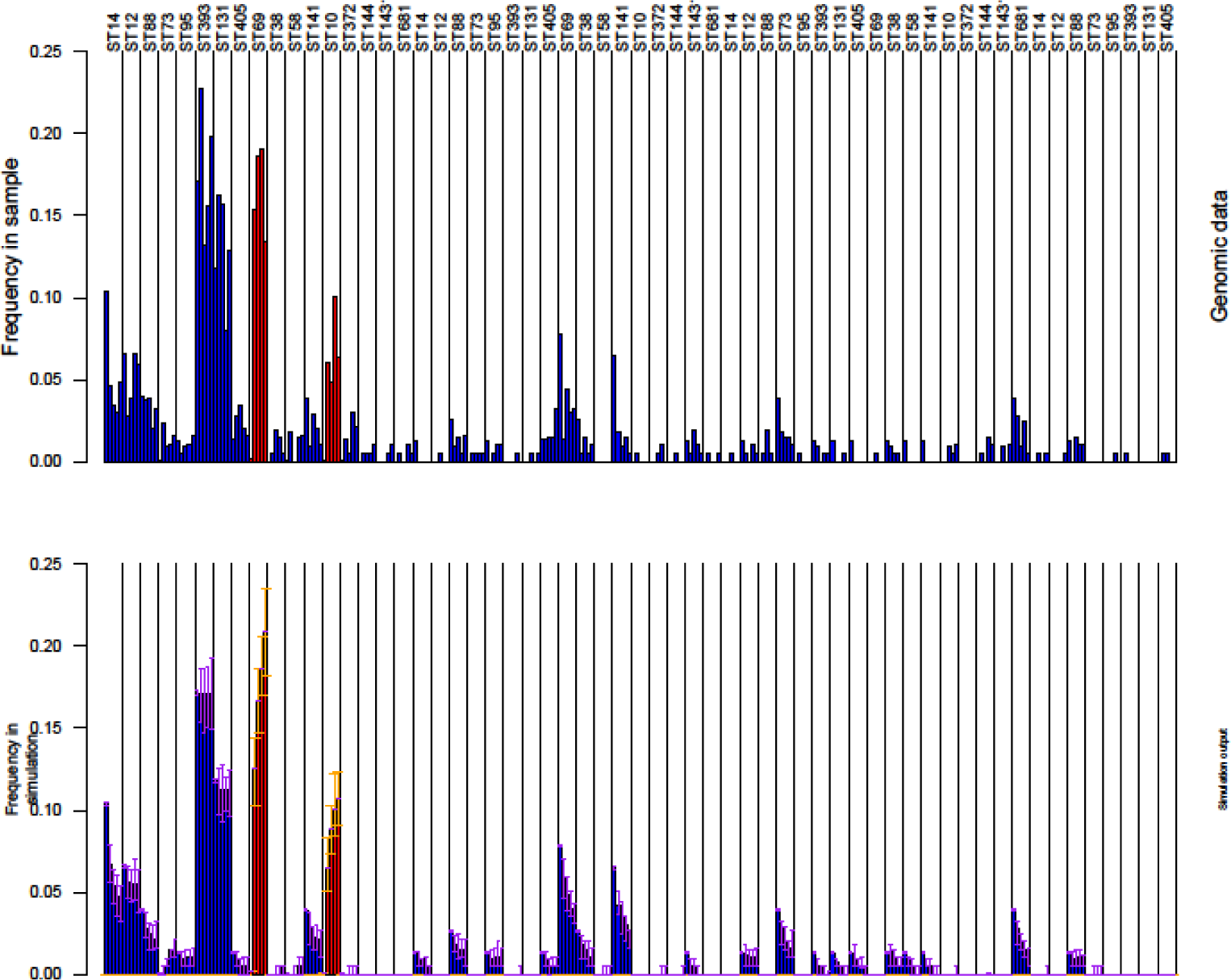
Full results of the NFDS simulations. These barcharts show the frequencies for all lineages from the one hundred simulations performed using the optimal parameters identified within the BOLFI model fitting, which are summarised in Fig 2. Each column again corresponds to a sequence cluster, and is annotated according to the predominant sequence type. The five bars within each column represent the frequency of the sequence cluster over subsequent time intervals: either that observed in the genomic samples for the top panel, or the median frequency in simulations in the bottom panel. The error bars on the bottom panel indicate the interquartile range for each bar from the 100 simulations. The red bars correspond to the ST69 and ST131 sequence clusters that had a reproductive fitness benefit, *r*, over the rest of the population.

**Figure S4:**
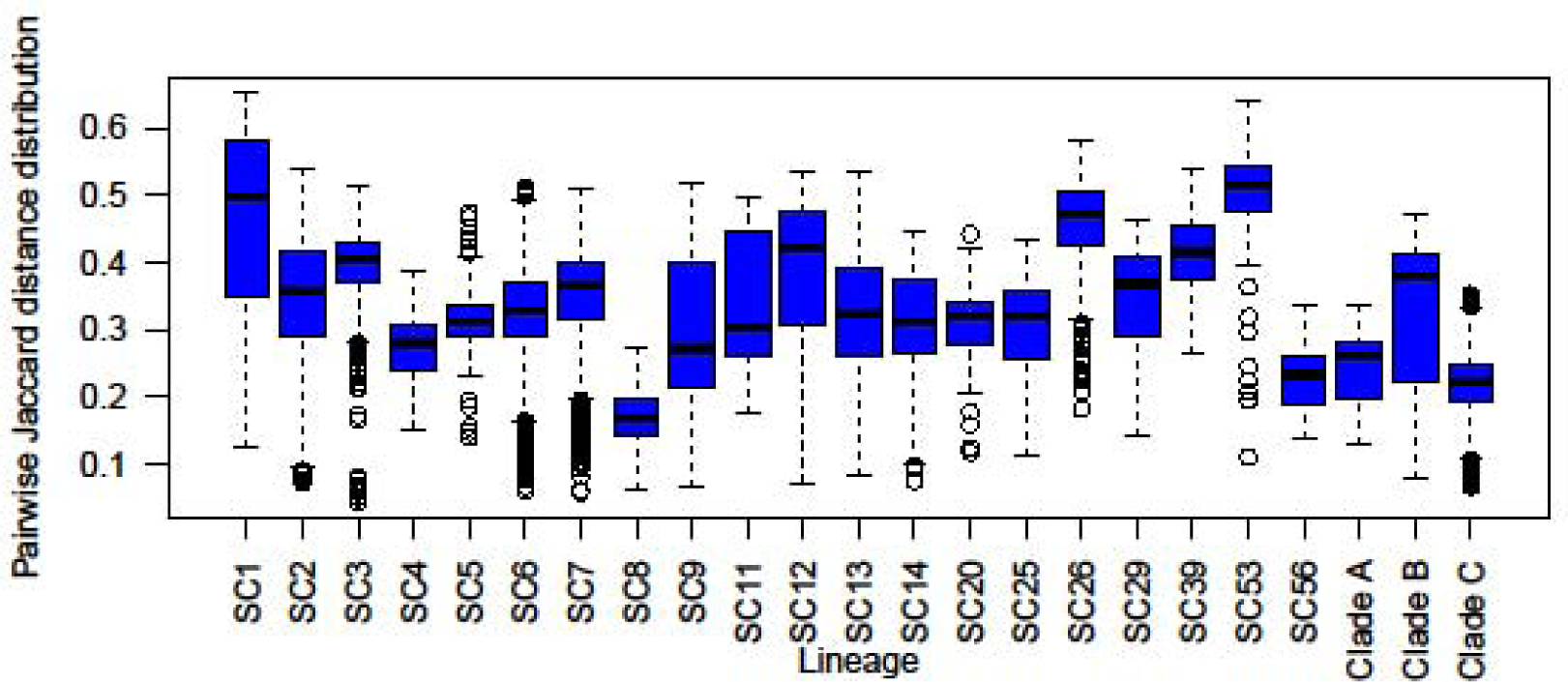
Diversity of intermediate frequency loci within *E. coli* lineages. The dissimilarity between pairs of isolates was measured as the binary Jaccard distance between them, based on the presence or absence of the intermediate frequency loci simulated in the multilocus NFDS model. The genetic diversity of each sequence cluster represented by at least ten isolates in the BSAC collection, and the three clades of the ST131 *E. coli*, are represented by a boxplot that shows the distribution of all such pairwise comparisons within the sequence cluster. This demonstrates the success of ST131 cannot be attributed to it exhibiting a greater diversity of loci under selection in the model relative to other lineages.

**Figure S5:**
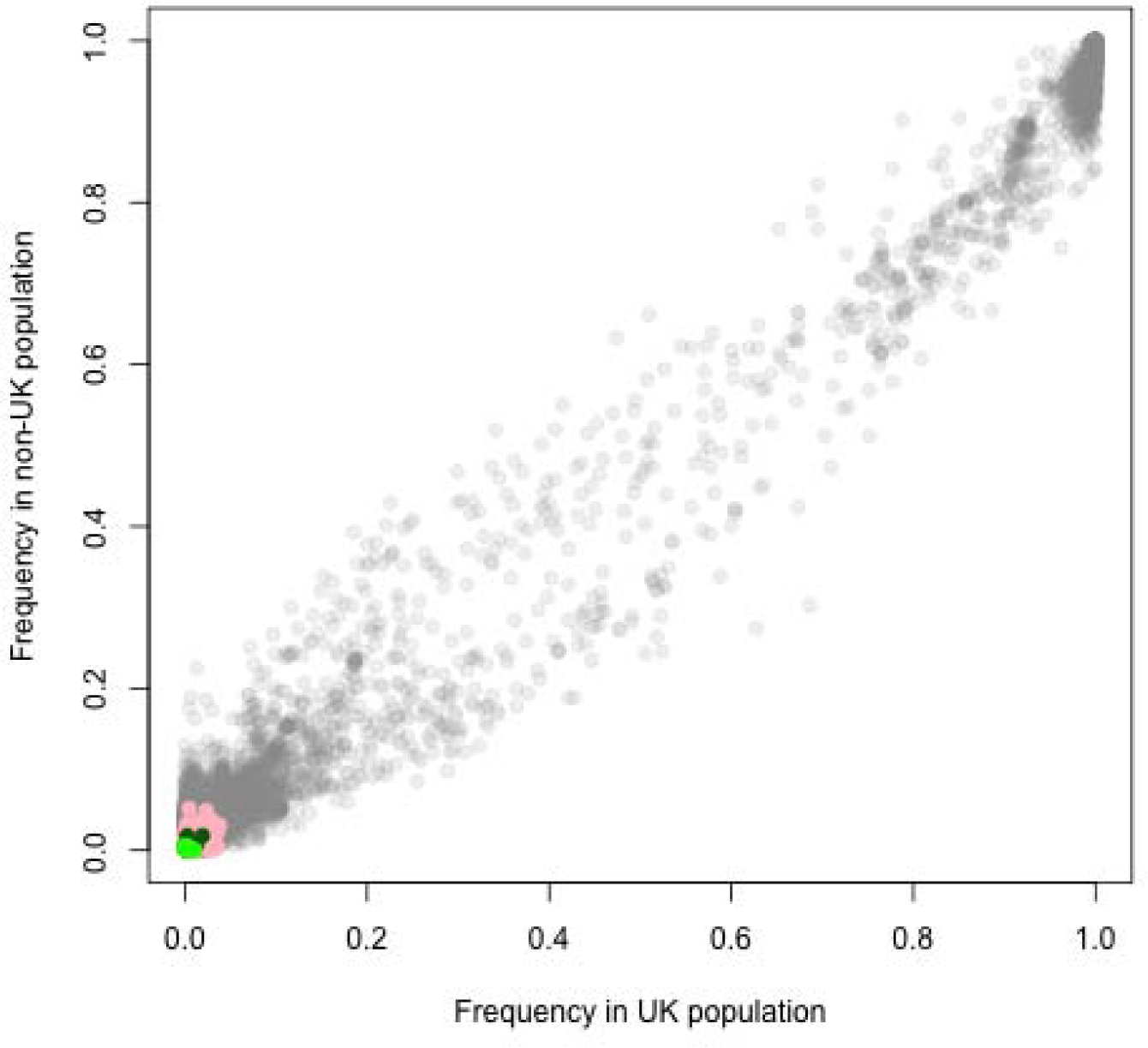
Frequency dependence plot showing the frequency at which all *E. coli* ST131 accessory genes occur in strains isolated from the UK versus strains isolated from outside the UK. The allele variants identified colour coded as in the previous figures: anaerobic metabolism (blue boxes), capsule production (pale blue boxes), cell division (black boxes), iron acquisition (orange boxes), pili/fimbriae production (green boxes), flagella (red boxes), and MDR efflux pumps (pink boxes)

**Figure S6:**
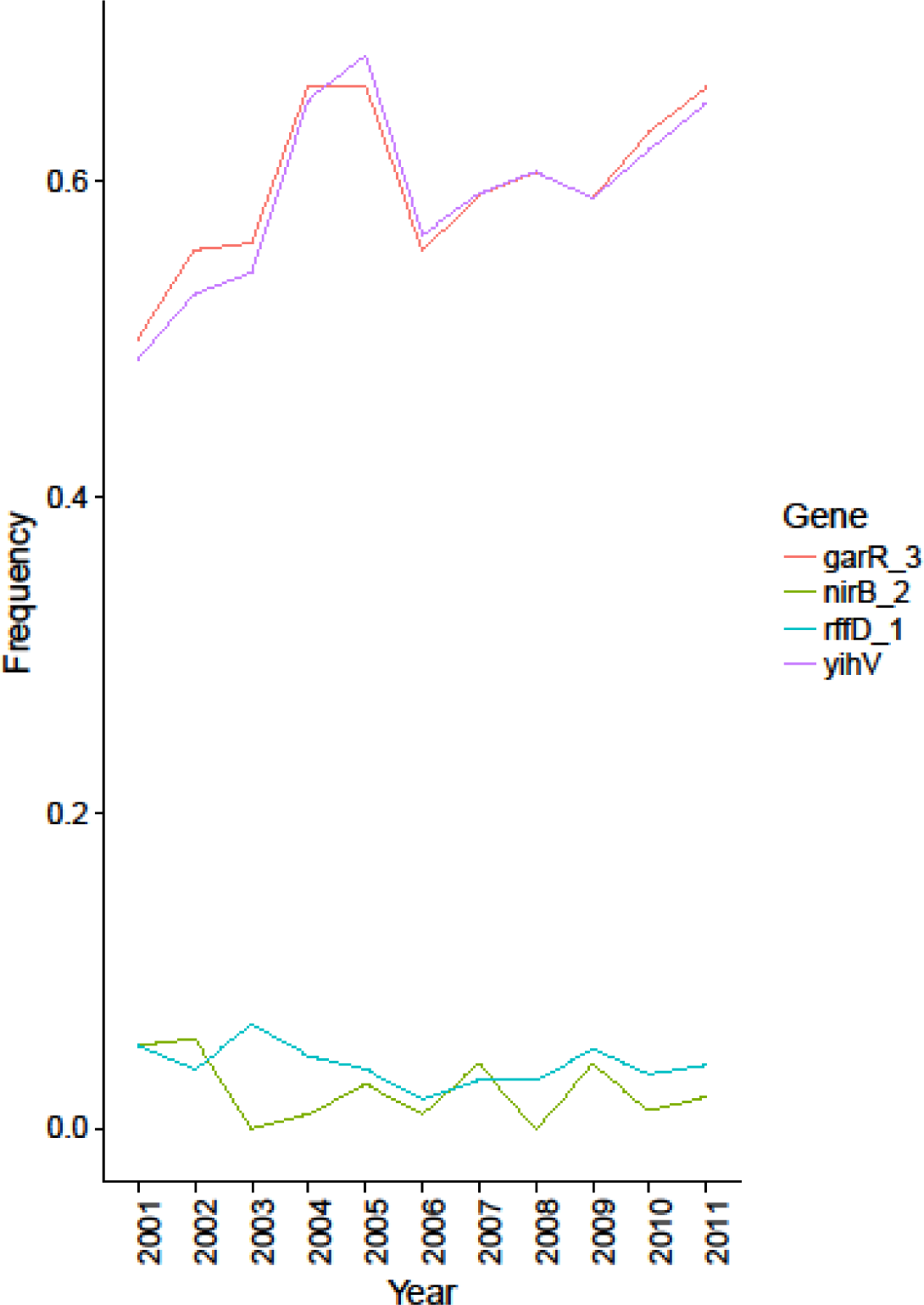
Stable intermediate frequencies of anaerobic metabolism loci. Four genes involved in anaerobic metabolism were found to be present at intermediate frequencies in the BSAC collection. All were absent from the ST131 lineage, except nirB_2, which was found in a subset of the lineage. Nevertheless, plotting their annual frequencies reveals distinct, stable frequencies over the period, despite the rise to prominence of ST131.

